# Low-frequency vibrational modes in G-quadruplexes reveal the mechanical properties of nucleic acids

**DOI:** 10.1101/2020.03.16.993873

**Authors:** M. González-Jiménez, G. Ramakrishnan, K. Wynne

## Abstract

Low-frequency vibrations play an essential role in biomolecular processes involving DNA such as gene expression, charge transfer, drug intercalation, and DNA–protein recognition. However, understanding of the vibrational basis of these mechanisms relies on theoretical models due to the lack of experimental evidence. Here we present the low-frequency vibrational spectra of G-quadruplexes (structures formed by four strands of DNA) and B-DNA characterized using femtosecond optical Kerr-effect spectroscopy. Contrary to expectation, we found that G-quadruplexes show several strongly underdamped delocalized phonon-like modes that have the potential to contribute to the biology of the DNA at the atomic level. In addition, G-quadruplexes present modes at a higher frequency than B-DNA demonstrating that changes in the stiffness of the molecule alter its gigahertz to terahertz vibrational profile. These results demonstrate that current theoretical models fail to predict basic properties of the vibrational modes of DNA.

**Statement of significance:** A number of recent studies have identified thermally excited low-frequency vibrational modes as a key deciding factor in the biological function of DNA. However, the nature of these vibrational modes has never been established. Here, vibrational spectroscopy with unrivalled signal-to-noise in the gigahertz to terahertz range is used to determine the low-frequency Raman spectra of nucleotides and oligomeric DNAs carefully chosen to form G-quadruplexes, structures formed by four strands of DNA common in the genome. These G-quadruplexes exhibit an unusual group of highly-underdamped delocalized vibrational modes—not reproduced by any of the theoretical models in use—which are expected to be the thermally excited. This provides a new perspective on the role of low-frequency vibrational modes in protein interactions and allostery.

## Introduction

The human genome contains in its transcriptional regulatory regions sequences rich in guanine that lose their double-helical structure in the presence of potassium cations to adopt a secondary conformation called a G-quadruplex (G4) (1, 2). This structure connects up to four strands of DNA (Fig. 1-B) forming a stack of planar tetramers of guanine (G-quartets, Fig. 1-A) linked together through Hoogsteen hydrogen bonds (Fig 1-C) (3). The potassium cations, which fit into the central space, stabilize the structure by their interaction with the inner carbonyl oxygen atoms of the guanine bases (4).

**Fig. 1.**
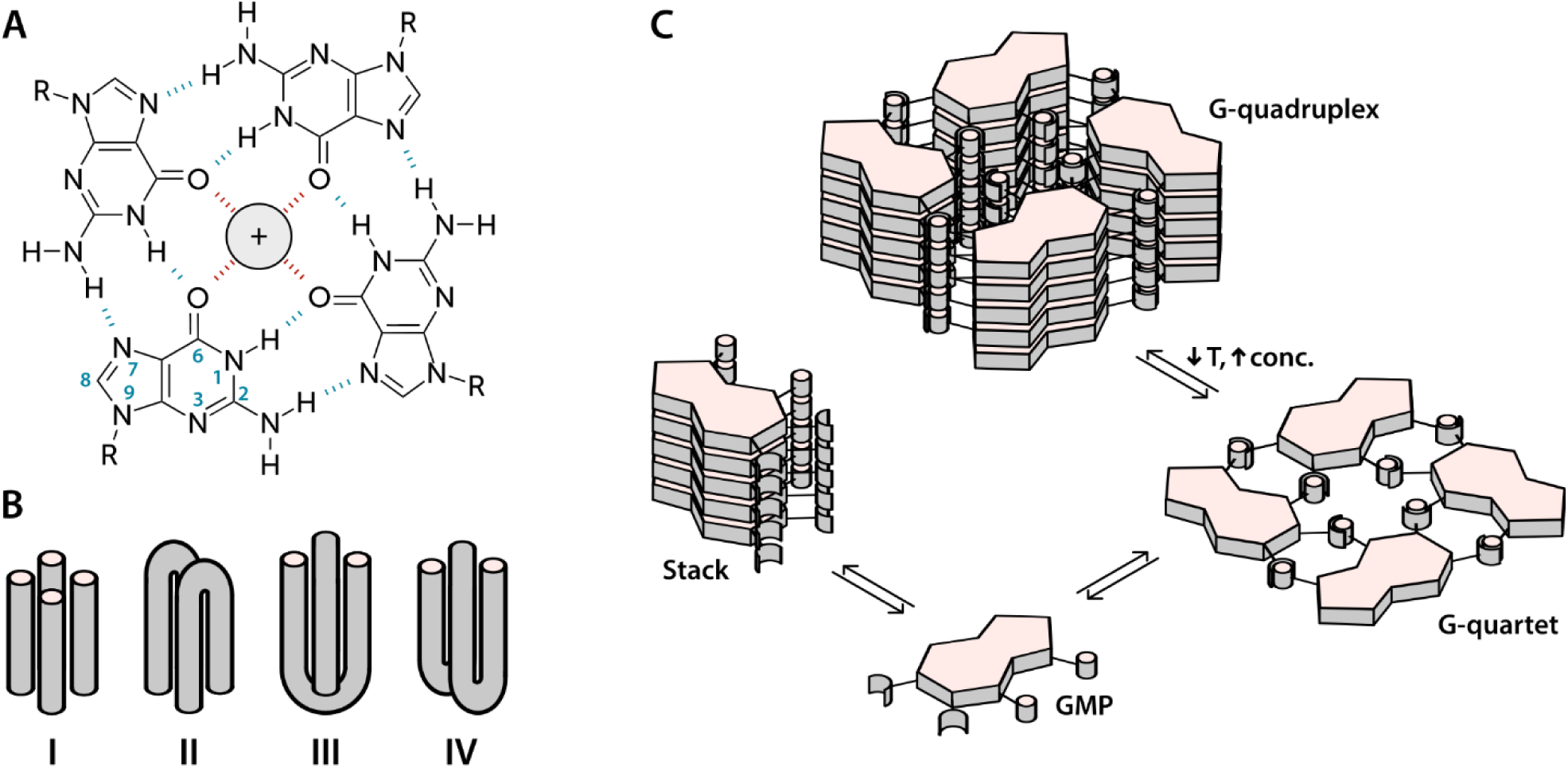
Structure and topology of G-quartets and G-quadruplexes. **(A)** Chemical structure of a G-quartet coordinated with a metal cation. Blue dashes represent Hoogsteen hydrogen bonds and red dashes the hydrogen-bonding interaction between the monovalent cation and the inner oxygen (O6) atoms. **(B)** Different topological variants of G-quadruplexes (G4s) depending on the number of strands: I - tetramolecular G4, II – bimolecular and parallel dimer of hairpins, Ill – bimolecular and antiparallel dimer of hairpins, and IV – intramolecular G4. **(C)** Equilibria involved in the self-association of guanosine monophosphate (GMP) in aqueous solution. The polar medium of the solvent promotes nucleotide stacking. Low temperatures and high concentrations contribute to the formation of G-quartets which are more stable in the G4 conformation.

There is mounting evidence that the alternative G4 structure plays an important role in the regulation of genes and oncogenes (5). Not only are G4 structures stable, but their prevalence is also controlled by the concentration of potassium ions, which is known to change with time and location within a cell (6). Furthermore, G4-promoting sequences are conserved through evolution (7) and there are a large number of proteins (Topo I, Pur1, MyoD, etc.) that have more affinity for G4s than for double-helix conformations (8). Particularly important amongst these proteins is the helicase protein encoded by the gene BLM, which is believed to undo G4s that are accidentally formed during recombination or replication. This gene is defective in Bloom’s syndrome, a rare genetic disorder characterized by genomic instability and a very high incidence of cancer (9).

G4 forming sequences are also abundant in the telomeres of chromosomes (1) and it has been suggested that these structures are formed between homologous chromosomes during meiosis (6). Furthermore, G4s inhibit the activity of telomerase—the enzyme that adds DNA to the ends of chromosomes to prevent them from shortening during cell division—that is often overexpressed in cancer cells (10).

The need for a better understanding of the role of non-canonical structures in genetic function and the potential of G4s as therapeutic targets for anticancer drug development has triggered extensive research.

The attention of the physical sciences to these conformations has focused mainly on the characterization of structures, their thermodynamic stability and their rate of folding and unfolding. (11–14) However, the dynamics of G4s have been almost completely ignored (except for some initial studies with NMR and 2DIR (15)) and their biological function is still unknown, despite the importance of low frequency vibrations in the biomolecular processes in which G4s play a role (16).

In this work, we present the first study of delocalized vibrational dynamics of G4s using femtosecond optical Kerr-effect (OKE) spectroscopy. OKE spectroscopy measures the low-frequency depolarized Raman spectrum in the time domain and is a technique that has proven to be critical in the investigation of the terahertz dynamics of DNA (17, 18) and other biomolecules in solution (19). We present the OKE spectra of 5’-guanosine monophosphate (GMP), a molecule that forms quartets and quadruplexes depending on the temperature and concentration (Fig. 1-C). At low temperatures and high concentrations, the G4 conformation is favored while at high temperatures and low concentrations, the equilibrium is shifted towards the formation of stacks (5). In order to disentangle the spectra of stacks, quartets, and G4s, we have compared the OKE spectra of GMP with the OKE spectra of solutions of the other DNA bases, adenosine, thymine, and cytosine monophosphate (AMP, TMP, and CMP), which cannot establish Hoogsteen bonds and just form stacks (20). We have also studied the effect of the size of the cations located at the central channel on the vibrational dynamics of G4s. While Li^+^ is too small to effectively stabilize the structure, Na^+^ is sized to be coordinated with four oxygen atoms in the G-quartet plane and K^+^ is sandwiched between quartets where it is coordinated with oxygens from both planes. Larger cations occupy the same position as potassium but distort the guanine quartets and stabilize the structure less. Finally, we have measured the OKE spectra of three DNA oligomers that form G4s with three different topological variants (see Fig. 1): a unimolecular G-quadruplex, a bimolecular complex formed from hairpin dimerization, and a parallel stranded tetraplex. We will show that these DNA conformations present a completely different vibrational dynamics from that of the single strands and double helix. This difference may be related to the biological function of the G4 structures, since the vibrational motions of the DNA influence the thermodynamics and mechanisms of the interaction between the nucleic acid and the proteins involved in genetic regulation.

## Materials and Methods

### Materials

Guanosine 5’-monophosphate disodium salt hydrate (from yeast, ≥ 99.9%, Sigma), adenosine 5’-monophosphate disodium salt (99.7%, Alfa Aesar), thymidine 5’-monophosphate disodium salt (99.7%, Alfa Aesar), lithium chloride (≥ 99%, Sigma), potassium chloride (≥ 99%, Sigma), and cesium chloride (≥ 99%, Sigma) were all used as received. Lyophilized, HSPF purified oligomers were prepared by Eurofins Genomics. HPLC gradient grade water from Fisher Chemical was used for preparing aqueous solutions. These solutions were filtered prior to the measurements using a 0.2 mm hydrophilic polytetrafluoroethylene (PTFE) filters (Millipore). DNA solutions were annealed by equilibrating them in a water bath at 363 K, followed by slow cooling down to the desired temperature.

### OKE experimental details

The OKE data were recorded in a standard time-domain step-scan pump-probe configuration and Fourier transformed to obtain the frequency-domain reduced depolarized Raman spectrum as described previously (17). A laser oscillator (Coherent Micra) provided ∼10 nJ pulses (0.8 W average power) with a nominal wavelength of 800 nm at a repetition rate of 82 MHz providing a 20-fs pulse width in the sample. Time-domain OKE traces were acquired by scanning with a step size of 5 fs up to a delay of 5 ps and thereafter logarithmically increased to a maximum delay of 1.2 ns. The sample was contained in a rectangular quartz cuvette (Starna, optical path : 1 mm) held in a brass block that was temperature-controlled with a precision of ± 0.5 K.

### NMR experimental details

Proton NMR spectra were recorded in D_2_O with a Bruker AVIII 500MHz Spect rometer.

### OKE data analysis

The OKE spectra consist of several broad overlapping bands that are analyzed through curve fitting to several analytical functions. At the lowest frequencies, one finds processes associated with diffusive orientational relaxation of the molecules. The diffusive orientational relaxation of the monomer aggregates and oligomers in solution has been fitted here with a Debye function

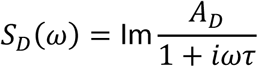

where *A*_*D*_ is the amplitude of the band, *ω* is the angular frequency, and *τ* is the relaxation time. The band at a slightly higher frequency, that can be assigned to the diffusive relaxation of water in the solvation shell of the solvents, cannot be fitted to a Debye function due to its much greater width. This band was modelled using a Cole-Cole function

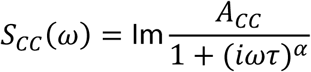

where *A*_*CC*_ is the amplitude of the band and *α* is a parameter that accounts for the broadness of the observed band.

In the terahertz range, one finds bands from modes that are not diffusive but critically damped or underdamped. These originate in librations, vibrations, and phonon-like modes. These have been fitted using the Brownian oscillator model (17)

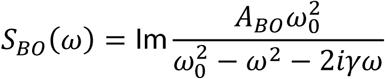

where *ω*_0_ is the undamped oscillator angular frequency and *γ* is the damping rate.

## Results

### The OKE spectra of GMP

The OKE spectra of disodium guanylate, Na_2_(5’-GMP), solutions were measured at temperatures between 283 and 358 K and concentrations between 0.1 M and 1.18 M (the solubility limit). To obtain the spectrum of solvated GMP, the contribution of bulk water was subtracted from the experimental OKE data (Note S1) and the resultant spectra are shown in Fig. 2. The low-frequency part of all spectra is dominated by a band peaking at frequencies too low to be accessible in these experiments and that moves to a higher frequency as the temperature increases. This band is characteristic of diffusive orientational and translational relaxation of GMP and could be modeled by the combination of a Cole-Cole and a Debye function (see OKE Data Analysis).

**Fig. 2.**
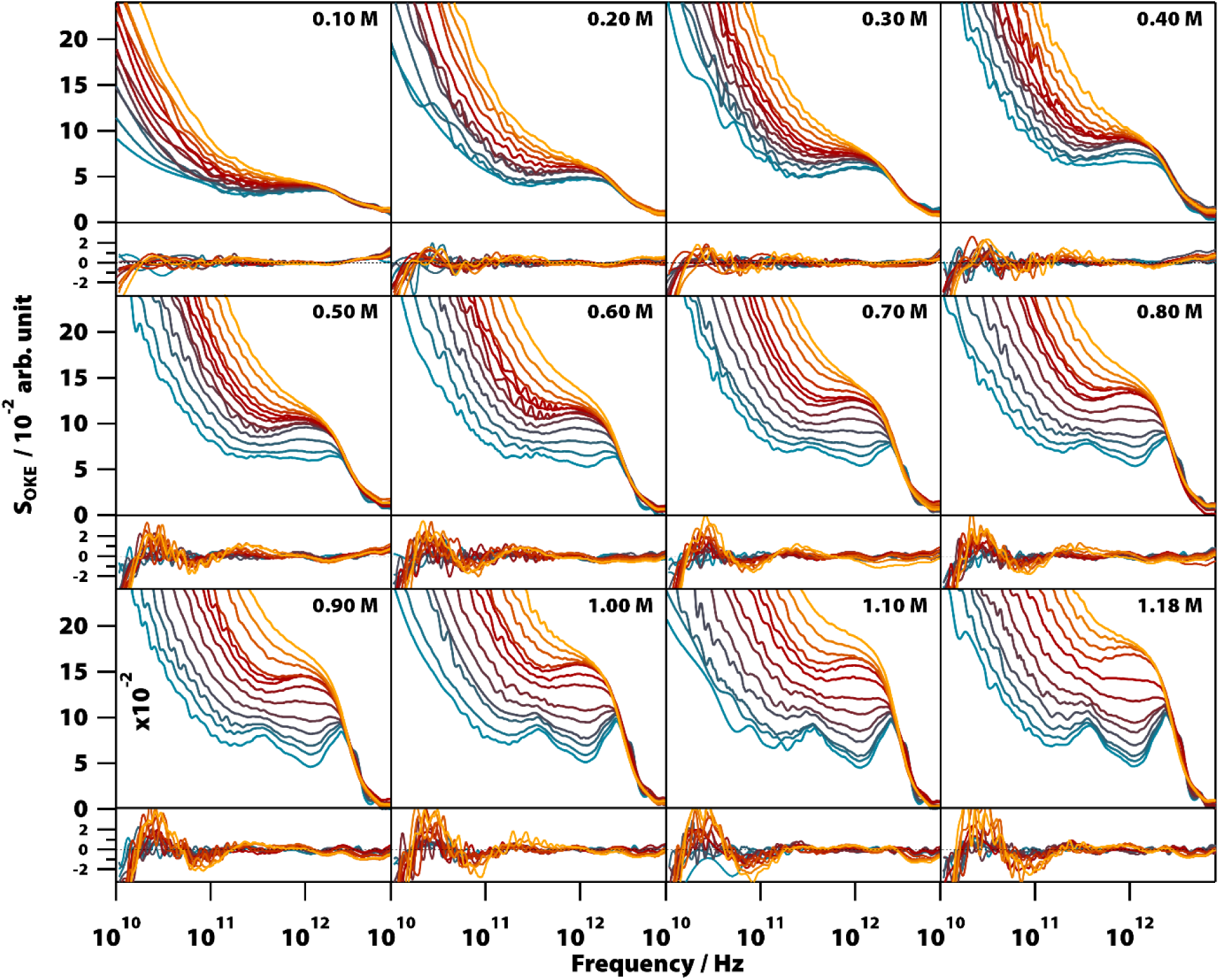
Optical Kerr-effect spectra of aqueous Na_2_(5’-GMP) solutions at different concentrations and temperatures showing changes in the spectra due to the formation of stacks and G-quadruplexes. Temperatures run from 283 (blue) to 328 K (red) in steps of 5 K and from 328 K to 358 K (orange) in steps of 10 K. The fitting deviations (residuals) between each spectrum and the proposed model values are shown in the graphs below the spectra.

The high-frequency part of the OKE spectra contained the most significant features with clear changes in shape as a function of temperature and concentration. At low temperature and high GMP concentration, the spectra show two clearly defined bands. The lower frequency band, peaking at ∼400 GHz, is unusually sharp for this frequency region. The higher frequency band in the region between approximately 1.5 and 4.5 THz is structured and appears to be the combination of three separate peaks. There is a weak band with a maximum at 5.5 THz that was sometimes difficult to observe due to the greater noise in the OKE spectra at high frequencies (19). On increasing the temperature or decreasing the GMP concentration, these sharp bands disappeared to leave the high-frequency region dominated by a wide band with a maximum at 1.6 THz that at higher temperatures fused with the low-frequency diffusive-relaxation band.

It was found that the high-frequency part of all the spectra shown in Figure 2 can be fitted accurately using a combination of two base spectra, each of these consisting of a number of Brownian oscillator functions (see Table S1). The first set, consisting of six Brownian oscillators (labeled B1-B6), fits the spectra of lower temperature and higher concentration, while the second set, consisting of two Brownian oscillators (labeled BA and BB). fits the opposite conditions. The relative amplitudes of the functions used in each set were nearly constant in all the spectra. It was found that the use of non-Lorentzian functions (i.e., Gaussians) did not allow a simpler, more accurate fit of all the 152 spectra shown in Fig. 2.

### OKE spectra of AMP and TMP

Aqueous solutions of these other bases at concentrations from 0.1 to 1.6 M were investigated to quantify the low-frequency vibrational modes associated with their structure (see Fig. 3; the OKE spectra for CMP were studied previously (17)). The OKE spectra of the solutions of all three molecules could be accurately fitted with two Debye functions for the low-frequency diffusive part and two slightly under-damped Brownian oscillators for the high-frequency part. Thus, the spectra of the AMP, TMP, and CMP solutions have an identical structure (same number of functions and the same approximate frequencies, damping rates, and relative amplitudes) to that of GMP at high temperature and low concentration. This demonstrates that the bands labeled BA and BB in GMP correspond to vibrational modes associated with the stacking interaction of nucleotides.

**Fig. 3.**
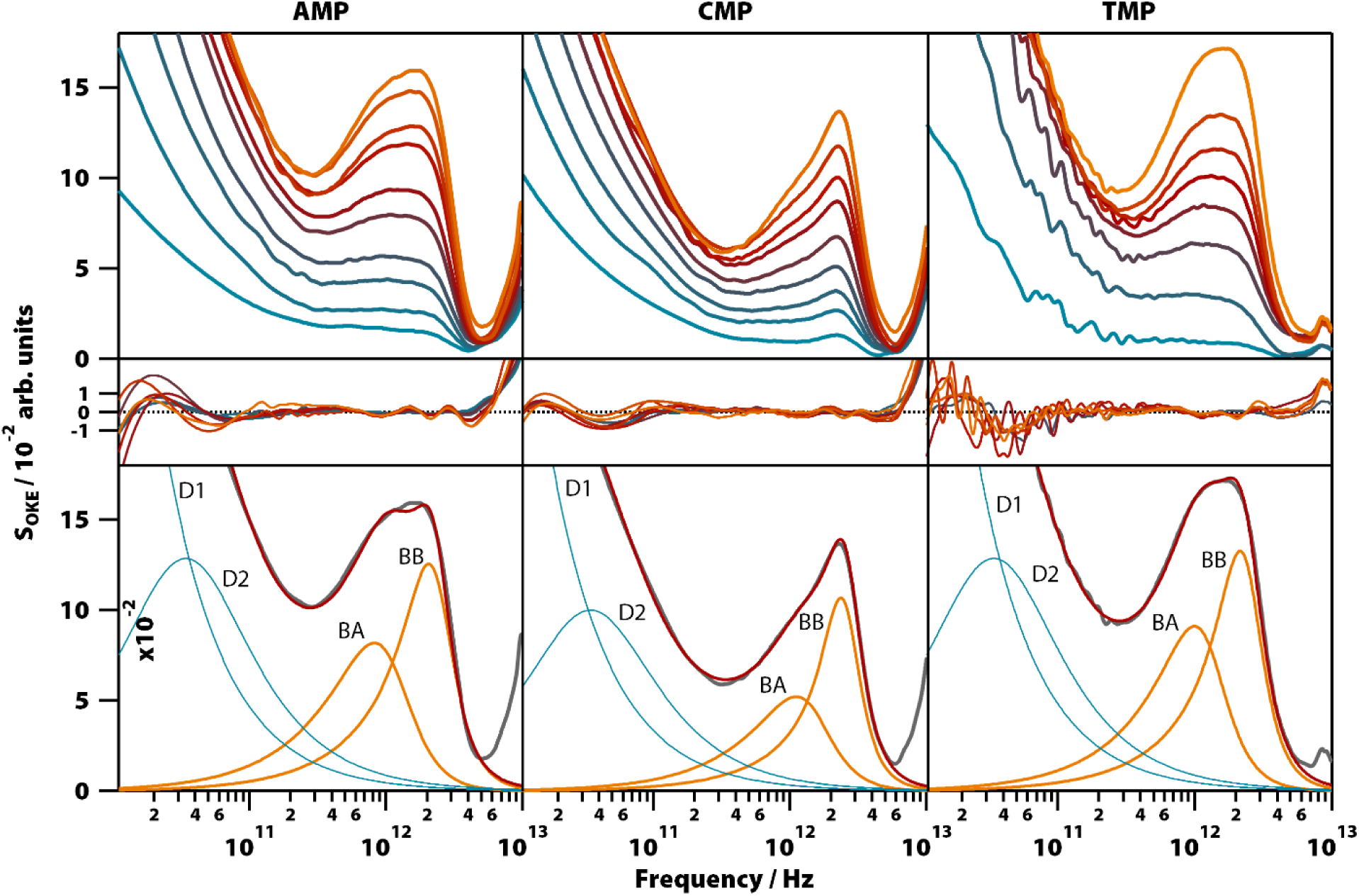
OKE spectra of aqueous solutions of three non-GMP nucleotides at 298 K demonstrating the fundamentally different form. Shown are the spectra for adenine (left column), cytosine (central), and thymidine monophosphate (right) at concentrations between 0.1 (blue) and 1.6 M (orange). Spectra are plotted in intervals of 0.1 M between 0.1 and 0.4 M and intervals of 0.2 M between 0.4 and 1.6 M for AMP and CMP and in intervals of 0.2 M for the whole concentration range of TMP (first row). The second row shows the fitting deviations (residuals) between each spectrum and the fittings and third row contains the 1.6 M spectrum for each nucleotide (grey), its fitting (red), and the functions used to model the data: Debye (blue) and Brownian oscillator (orange).

### ^1^H NMR spectra of GMP

Since the temperature- and concentration-dependent OKE spectra of GMP can be fitted with two base spectra, and since one of these base spectra has now been assigned to stacked GMP, it implies that the other base spectrum corresponds to the G4 structure (see Fig. 1). This premise was confirmed by measuring the ^1^H NMR spectra of a 1.18 M Na_2_(5’-GM P) solution between 283 and 353 K in intervals of 10 K (Fig. S1). At higher temperatures, these spectra show a sharp line around 8 ppm corresponding to the hydrogen 8 (see Fig. 1) of the guanine ring of unassociated GMP (4). As the temperature dropped, the intensity of this peak decreases while four peaks attributed to the G4 structure appear between 6.2 and 8.2 ppm (21, 22).

### GMP Equilibria and Thermodynamics

Averaging the intensity of the sets of bands in each base spectrum used to model the OKE data allowed us to determine precisely the proportion of the two conformations at each temperature and concentration (Fig. S2). From the dependence on temperature of the signal strength of the G4 structures, it is possible to determine their melting or denaturation temperatures for each initial concentration of GMP (Fig. S3 and Table S2). The consistency of our calculated values with the NMR spectra and with the melting temperatures found in the literature (22, 23) also support our assignment of each base spectrum. Employing these melting temperatures, we could calculate a standard-state van’t Hoff enthalpy and entropy changes of ΔH^0^ = 73 ± 6 kJ mol^−1^ and ΔS^0^ = 180 ± 20 J K^−1^ mol^−1^ for the dissociation of the G-quadruplexes (Fig. S3 and Note S2).

Fig. 4 shows the influence of the initial concentration of GMP on the intensity of each base spectrum for each temperature. These results show that the measured GMP solutions follow the chemical equilibria shown in Fig. 1-C. As the concentration increases, the signal of the GMP stacks increases until the formation of G4 structures is favored and begins to descend. In the case of the signal associated with G-quadruplexes, it rises linearly, as much as the temperature allows, as the GMP concentration increases.

**Fig. 4.**
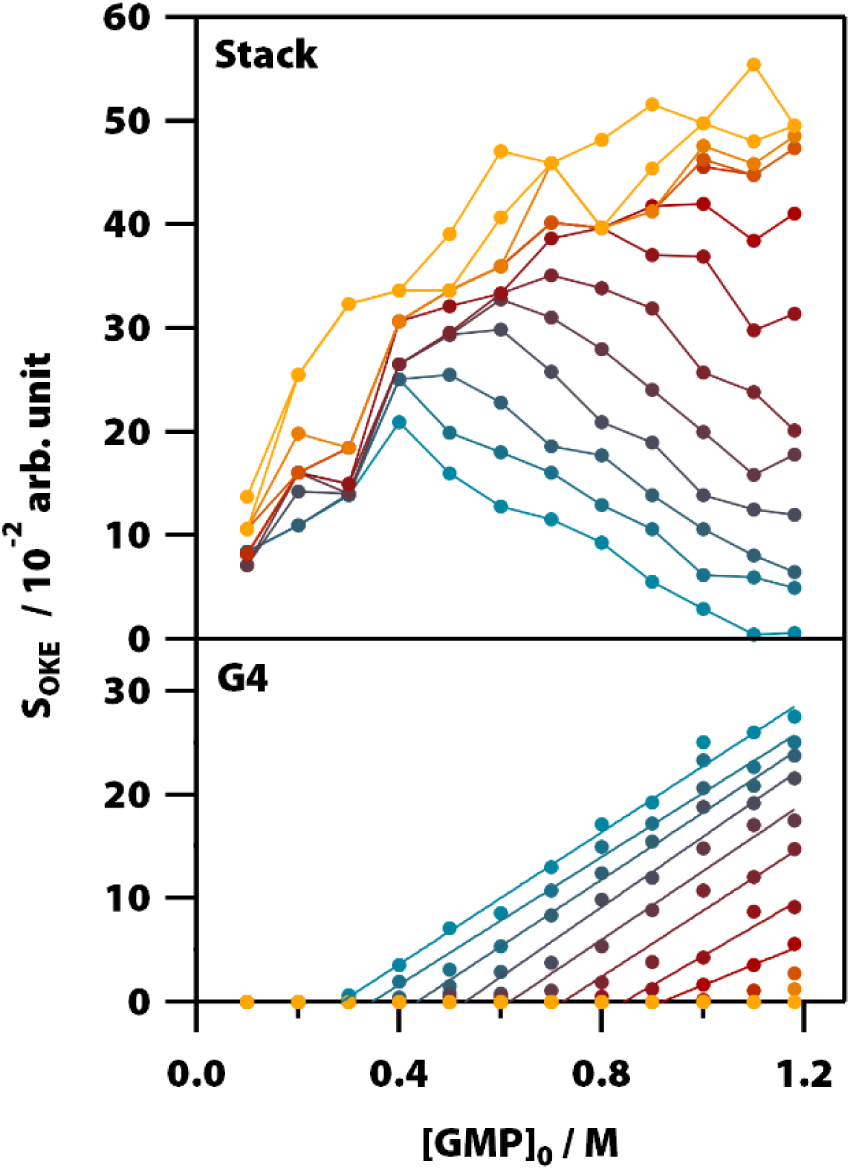
Influence of temperature and initial concentration of GMP on the averaged OKE intensity of the bands associated with the stack (top) and G-quadruplex (bottom) conformations showing the delayed onset of the latter. Temperatures run between 283 (blue) to 318K (red) in steps of 5 K and from 318 K to 358 (orange) in steps of 10 K.

### Influence of Cations

To gain more insight into the vibrational modes detected in the G4 structure, we compared in a broad range of temperatures (283 − 358 K) the OKE spectra of GMP·Na^+^ with the OKE spectra of three GMP solutions prepared with an excess concentration of a different monovalent cation: lithium, potassium, and cesium. These cations replace sodium in the inner channel of the structure, altering its structure, stability, and dynamics due to their size difference (24). As a result, the solubility of GMP also changes, forcing the concentration of the measured solutions to be different. The spectra obtained (Fig. 5, first row) could be modeled over the entire temperature span using the same number of Brownian oscillator functions employed in GMP·Na^+^ solutions.

**Fig. 5.**
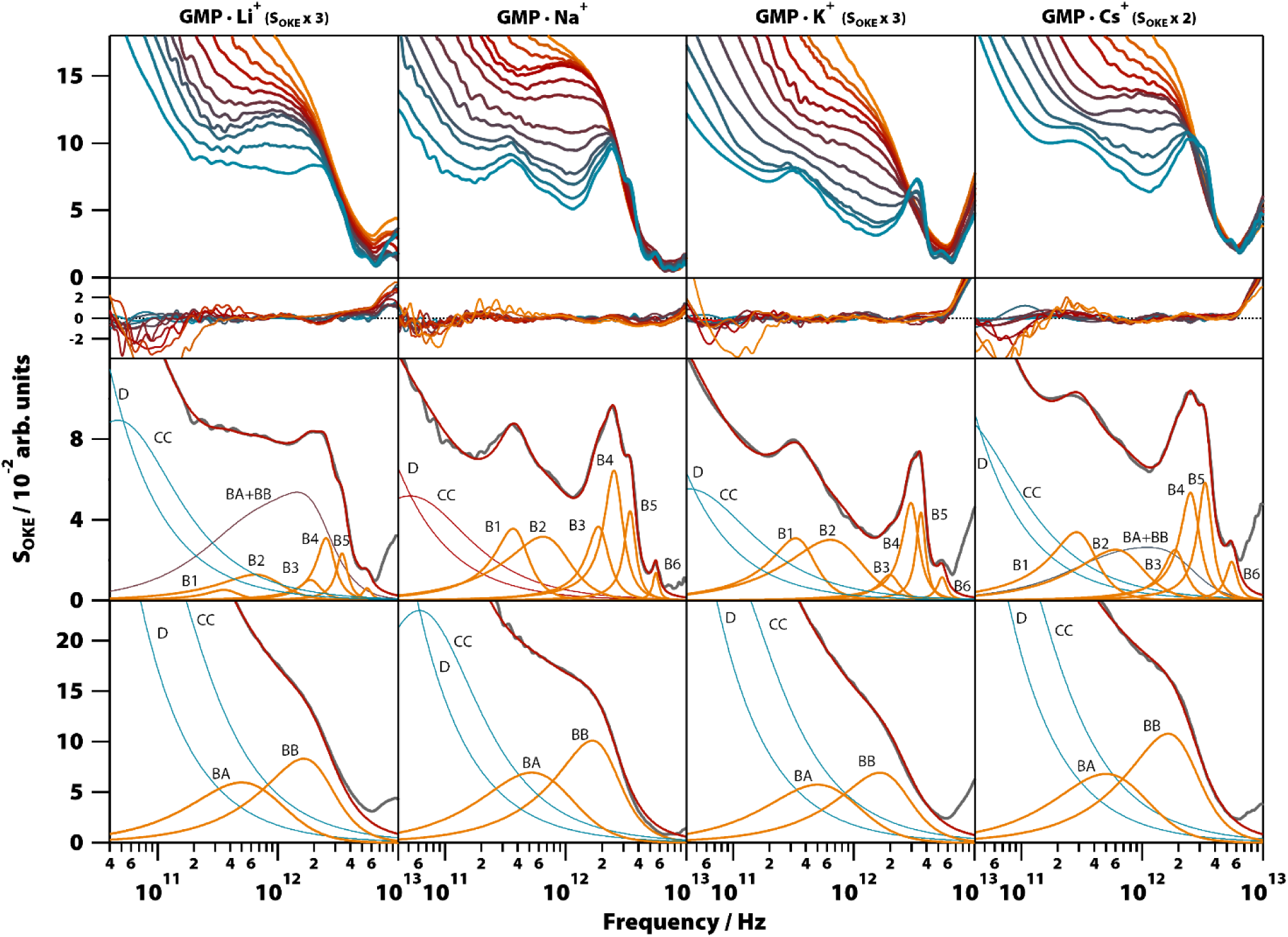
Temperature-dependent OKE spectra of GMP solutions with different cations. Top row, from left to right: OKE spectra of GMP in the presence of Li^+^ (GMP 0.25 M, LiCI 1.0 M), Na^+^ (Na_2_(5’-GMP) 1.0 M), K^+^ (GMP 0.2 M, KCI 0.5M), and Cs^+^ solutions (GMP 0.4 M, CsCI 1.0 M). The temperature runs from 283 (blue) to 328 K (red) in steps of 5 K and from 328 K to 358 (orange) in steps of 10 K. Below the residuals between each spectrum and the fittings are shown. The remaining rows contain the spectrum for each case (grey) at 283 K (third row) and 358 K (fourth row) and the functions employed to fit the spectra. Diffusion associated functions (Debye and Cole-Cole) are colored in red and vibration associated functions (Brownian oscillators) in orange. The blue band in the GMP·Li^+^ and GMP·Cs^+^ spectra at 283 K is the result of the sum of the two Brownian oscillators associated with GMP stack conformation (BA + BB).

Moreover, the fit parameters for the spectra at higher temperatures (the functions labeled BA and BB in Fig. 5, third row) were equal, supporting the assignment to vibrations in a stack of nucleotides, since the cations of the medium do not take part in these structures. At low temperatures, the cation does influence the OKE spectra associated with G4s. In lithium and cesium solutions it is observed that the spectra, even at the low temperature of 283 K, consist of the combination of the base spectra of both G4 and stacked GMP. This agrees with the fact that these cations are the ones that stabilize the structure the least; lithium because it is small in comparison with the central space and cesium because its size disrupts the structure. The characteristics of the bands are also affected by the size of the central cation. The bands of cations that are smaller than the central space (approximate radius: 131 pm, PDB entry: 1K8P), lithium (effective radius: 60 pm) and sodium (95 pm) (25), have essentially the same oscillatory parameters (Fig. 5, second row, and Table S1). However, the potassium cation, which is practically the size of the central space (radius: 133 pm), presents a spectrum very similar to that of sodium and lithium, except that its B4 band is displaced to a higher frequency position. The base spectrum when cesium is present (radius: 169 pm) has B1, B2 and B5 bands shifted to lower frequency positions, compared to the sodium spectrum.

### OKE spectra of oligonucleotide G4s

In order to confirm that the narrow spectral band shapes seen in G4s constructed from GMP nucleotides are more general, aqueous solutions of three rich-in-guanine, G4-forming DNA oligomers were studied: thrombin-binding aptamer d(GGTTGGTGTGGTTGG) (TBA), which forms an unimolecular G-quadruplex, d(G^3^CT^4^G^3^C), which forms a dimer of two parallel hairpins, and d(TGGGGT), which forms a parallel-stranded tetraplex. The spectra from these oligomers are compared with that of a double-helix forming oligomer, d(T_2_AT_2_A_3_TATAT_3_A_2_TA_2_), studied previously in the conformation of B-DNA (18).

Unlike GMP, these oligomers required the presence of K^+^ to form the G4 structure (Fig. S5), so the solutions of the oligomers were prepared with a concentration of 1.65 M of KCI (except TBA which formed a gel with that concentration and required KCI 0.4 M). The temperature-dependent OKE spectra (283 to 358 K) are shown in Fig. 6. As with GMP, the high-frequency part of these spectra can be fitted with two base spectra associated with a G4 structure and its denatured conformation.

**Fig. 6.**
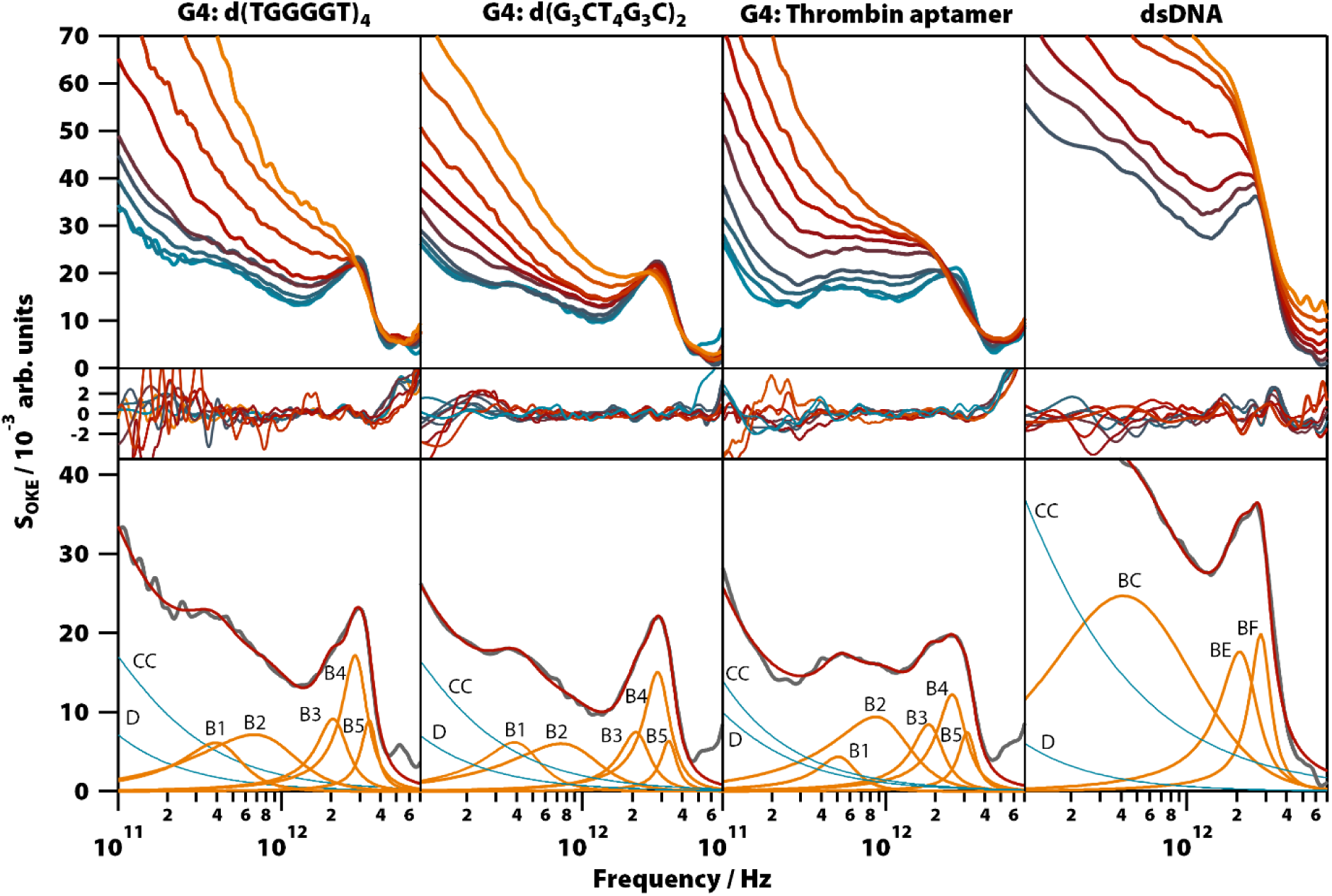
OKE spectra of solutions of three G4-forming oligomers and one double-helix forming oligomer demonstrating that the gigahertz-to-terahertz modes seen in GMP-based G-quadruplexes persist in oligomers. From left to right: Spectra of d(TGGGGT) (80 mg/ml, KCI 1.6 M), d(G_3_CT_4_G_3_C) (80 mg/ml, KCI 1.6 M), and TBA (80 mg/ml, KCI 0.4 M) between 283 (blue) to 318K (red) in steps of 5 K and from 318 K to 358 (orange) in steps of 10 K. Spectra of d(T_2_AT_2_A_3_TATAT_3_A_2_TA_2_)_2_ (right) between 298 (blue) and 358 K (orange) in intervals of 10 K. Residuals are shown below. The third row shows the spectrum of lower temperature for each oligomer (grey) and the functions used to model them. Debye and Cole-Cole functions are colored in red and Brownian oscillators in orange.

The base spectrum of the G4 conformation is again observed at the lowest temperatures and, for both oligomers, it can be adjusted with the same set of functions as used for the GMP solutions. Furthermore, the parameters of these Brownian oscillators (Table S1) deviate likewise from the models from G4 structures in GMP, suggesting that these vibrational modes have characteristics that appear only when nucleotides are bonded in oligomers. A comparison of these results with the model used to fit the double-helix spectrum (17) (Fig. 6, right column) showed that bands B3 and B4 coincide at the same frequency with the bands BE and BF of the double-helix model, respectively.

The bands of the three oligomers studied at high temperatures after denaturation (Table S1) are roughly the same and coincide with those of the single-stranded DNA model measured previously (17).

### Summary of results

Here we have shown that the terahertz-frequency vibrational spectra of Na_2_(5’-GMP) in aqueous solution are generated by the combination of the base spectra from two different structures whose proportion depends on their concentration and temperature. It was demonstrated that the predominant base spectrum at high temperature or low concentration is due to the vibrational modes associated with the stacked nucleotide structure. This base spectrum, which is the only one observed in solutions of other nucleotides that only stack, could be modeled using two slightly underdamped Brownian oscillators.

It was demonstrated that the second base spectrum is due to vibrations associated with the G-quadruplex (G4) conformation and could be observed in G4s consisting of GMP nucleotides as well as three exemplar G4-forming oligonucleotides. This demonstrates that this base spectrum is largely characteristic of the G4 structure and not very much dependent on the exact base sequence. These associations between spectra and structures allow one to determine the stability of G4s from the ratio between the intensities of both base spectra and to determine the effect that factors such as temperature or the size of the counterion have on that stability. Thus, it was confirmed that the observed structures behaved as expected, except in the experiments with GMP and Li^+^, whose spectra indicated that this cation does not destabilize the G4 structures, as suggested recently (25).

The G4-associated base spectrum can be fitted using six underdamped oscillator functions with center frequencies in the 0.4-5.5 THz range. Apart from the lowest-frequency band B1, the observed modes have a similar damping rate of around 0.5 THz. Therefore, the viscous drag of the hydration shell that dissipates the energy stored in the oscillations is less efficient as the frequency of the bands increases, enabling coherent vibrations that can contribute to the biological function of DNA at the atomic level (26).

The experiments presented here provide more information about the vibrational modes detected in G4s. The band B1 is highly unusual with a peak frequency of only ∼400 GHz and a damping rate of 150-190 GHz making it highly underdamped. In all liquids, solutions, and biomolecules studied in this frequency region, an underdamped mode below 1 THz has never been observed except in vitrified solutions (27, 28). Using a speed of sound of 3 km/s as measured in DNA (29), the 400-GHz mode could correspond to an acoustic standing wave fitting in a phononic cavity of length 3.75 nm, which is the approximate circumference of a G-quartet. This interpretation is supported by the shift to lower frequency exhibited by this band when the inner cation is Cs^+^. As shown by structures characterized by x-ray crystallography (PDB entries: 2GW0, 1K8P and 1JB7), cations larger than potassium distort the guanine quartets, increasing the space between the molecules and, consequently, the length of the phononic cavity. The band B2 peaks at approximately double the frequency of the B1 band and has the same behavior towards the cations of the inner channel, suggesting that this might be an overtone.

The properties of the band B3 in the solutions of GMP are independent of the coordinating cation. However, the band is shifted to a higher frequency in the studied oligomers. Therefore, this mode is influenced by the restriction in the movement caused by the bonding between monomers. The fact that bands B3 (in G4-forming GMP and oligonucleotides), BE (in melted G4-forming oligonucleotides), and BE (in other DNA oligonucleotides, either single-stranded or double-stranded) all have similar vibrational parameters at the same frequency implies that the bases of the nucleotides do not participate in this vibrational mode, but are due to the interactions between the sugar and phosphate groups.

The band B4 is an underdamped vibrational mode that shifts to a higher frequency when potassium, the cation that most stabilizes the structure, is present. The band B4 in the oligomers d(TGGGGT)_4_ and d(G_3_CT_4_G_3_C)_2_ which were also measured in solutions with KCI, have characteristics almost identical to the spectra of the GMP-K^+^ complex. This dependence of K^+^ excludes that, although they appear in the same frequency and have a similar damping ratio, the B4 band in the oligomers and the BF band observed in the double helix of the DNA are related.

The bands B5 and B6 are two highly underdamped vibrational modes that appear exclusively in the OKE spectra of nucleic acids in the G4 conformation. Their parameters are roughly equal in all the compounds studied here.

## Discussion

The low-frequency dynamics of DNA can be modeled theoretically using simple ball and spring models (30), coarse-grained models (31), and atomistic molecular models (32). Such models are too crude to reproduce the G4 and B-DNA spectra presented here. However, coarse-grained models analyzed using Instantaneous Normal Modes (INM) show that terahertz-frequency modes in DNA can be understood in terms of delocalized phonon-like modes in an approximately one-dimensional structure, localized by disorder and interactions with the surrounding solvent (33). In a perfect (ordered, undamped, and infinite) lattice, Raman scattering (including OKE) should only occur from optical phonons at the zone center (that is with a wavelength delocalized over the entire structure). However, in the presence of disorder and damping, momentum selection rules are expected to break down and Raman-scattering will take place from essentially all optical and acoustic phonon modes. Previous simulations (33) have suggested that this should lead to a broad 2-3 THz band but that is not seen in these experiments. Similarly, normal-mode analysis applied to different nucleic acids (34–37) results in a relatively homogeneous distribution of low-frequency modes up 5 to 8 THz (150-250 cm^-1^) in which the modes with frequencies > 1 THz have the greatest contribution to the motion of the atoms of the molecule. Thus, current vibrational models of DNA singularly fail to predict the small number of distinct vibrational bands observed in this work. This calls into question the validity of such models for calculating thermodynamic properties associated with DNA folding and DNA-protein interactions.

It is usually assumed that terahertz frequency modes in molecules in aqueous solution will be strongly overdamped (38). However, the data presented here show that this is not the case: double-stranded DNA has modes at ∼2 THz that are underdamped (damping rate 0.5-0.7 THz) while the G4 structure has modes at even higher frequencies (approximately 2, 2.6, 3.4, and 5.5 THz). As can be seen in Table S1, the damping of the modes labeled BA and BB in nucleotide stacks is approximately 1 THz, whereas that of the modes labeled B1-B6 in the G4 structures ranges from 0.15 to 0.5 THz. This reduced damping reflects the greater rigidity of the G4 structure over dsDNA and reduced coupling to the solvent. As *k*_*B*_*T/h* at room temperature is 6.1 THz, it is these modes that play the most important role in determining the vibrational enthalpy and entropy.

Long-distance signal transmission and allostery have been demonstrated in model systems such as the complex between DNA and two DNA binding proteins (a eukaryotic transcription factor and a type II endonuclease (39)). It has been suggested that the oscillatory distance dependence of the allosteric interaction mediated by DNA is associated with thermally excited low-frequency vibrational modes of duplex DNA (dsDNA). This allostery is largely dependent on changes in mechanical properties of the linker DNA: as the interaction with the first DNA-binding protein stiffens the DNA, the entropic penalty for binding the second DNA protein is less resulting in allostery.

A similar effect may be at play in the G4 structures. Nucleotide G4s have phonon bands at a higher frequency than in dsDNA. This will result in a free energy penalty that is compensated for by the interaction with the central cation. However, it also causes the G4-structure to be much stiffer than dsDNA and hence binding to it will incur a much-reduced entropic penalty. This effect was also observed in G4 structures formed by GMP when the inner cation was changed. The structures most stabilized by the inner cation, and therefore the stiffest, are the structures that show phonon bands at a higher frequency. Likewise, the bands observed in the G4 structures with their nucleotides linked by phosphodiester bonds, d(TGGGGT)_4_, TBA, and d(G_3_CT_4_G_3_C)_2_, appear at a higher frequency than the bands of G4 structures formed by free GMP molecules. All these experiments reinforce the idea that phonon-like modes play an important role in allosteric interaction and, as a result, in gene regulation.

## Author contributions

All authors contributed to the study and manuscript. M.G.-J. and G.R. were responsible for data collection and numerical anal ysis. M.G.-J was responsible for sample preparation and basic characterization. M.G.-J. and K.W. carried out data analysis and manuscript preparation.

## Competing interests

The authors declare no competing financial intere sts.

## Data and materials availability

The data that support the findings of this study are available in Enlighten: Research Data Repository (University of Glasgow) with the identifier: http://dx.doi.org/@@@.

## Acknowledgments

We thank the Engineering and Physical Sciences Research Council (EPSRC) for support through grants EP/J009733/1, EP/K034995/1, EP/N508792/1, and EP/N007417/1. This work was partially funded by Leverhulme Trust Research Project Grant RPG-2018-350.

